# Origins of fractional control in regulated cell death

**DOI:** 10.1101/201160

**Authors:** Luís C. Santos, Robert Vogel, Jerry E. Chipuk, Marc R. Birtwistle, Gustavo Stolovitzky, Pablo Meyer

## Abstract

Individual cells in clonal populations often respond differently to environmental changes; for binary phenotypes, such as cell death, this can be measured as a fractional response. These types of responses have been attributed to cell-intrinsic stochastic processes and variable abundances of biochemical constituents, such as proteins, but the influence of organelles has yet to be determined. We use the response to TNF-related apoptosis inducing ligand (TRAIL) and a new statistical framework for determining parameter influence on cell-to-cell variability through the inference of variance explained, DEPICTIVE, to demonstrate that variable mitochondria abundance correlates with cell survival and determines the fractional cell death response. By quantitative data analysis and modeling we attribute this effect to variable effective concentrations at the mitochondria surface of the pro-apoptotic protein Bax. Further, we demonstrate that inhibitors of antiapoptotic Bcl-2 family proteins, used in cancer treatment, may increase the diversity of cellular responses, enhancing resistance to treatment.

---

Isogenic populations of cells in homogeneous environments have the seemingly paradoxical capacity to generate many unique cell states. This ability is found in many, if not all, types of single celled organisms and in the distinct cell types of multicellular organisms. For example, *B. subtilis* cells were shown to independently and transiently switch between vegetative and competent states [1], hematopoietic progenitor cells can differentiate into either erythroid or myeloid lineages [2], and cancerous tissue maintain distinct sub-populations throughout the course of disease [3]. In all such cases, a cell’s propensity for a particular state is attributed to the intrinsic stochasticity of low-copy number biomolecular reactions [4–6], or extrinsic variations in the abundances of its components [7–9]. Taken together it is clear that stochastic transitions of cell state that are driven by non-genetic sources of cell-to-cell variability (CCV) are fundamental to the maintenance of single cell populations, the function of distinct tissues, and structure of clinical lesions in diseases such as cancer.

One commonly studied source of CCV is protein abundance. Its premier status as a dominant source of non-genetic CCV is due to its stochastic production [6, 10], and the sensitivity of cellular decision-making machinery to variations in their components. For example, in biological signal transduction, information regarding the cell’s environment is processed by a cascade of biomolecular reactions. Variation from one cell to another in any one of the corresponding biomolecules varies the signal magnitude across the population, making unique the cell’s perception of environmental conditions and its corresponding response [11–14]. While it has been definitively shown that CCV in protein abundance influences cellular decisions, little attention has been given to other non-genetic sources of CCV.

There are numerous examples in which non-genetic and non-protein sources of CCV are conjectured to impact biological phenomena. For example, centrosome abundance [15], the size of the Golgi apparatus [16], and mitochondria abundance [17–20] all have been shown to vary from cell-to-cell. To determine if diversity in cell behaviors may be attributed to CCV in organelle abundance, our study focuses on the role of mitochondria in the context of TRAIL induced apoptosis.

Indeed, the abundance of mitochondria per cell has been shown to positively correlate with a cell’s propensity for apoptosis [20]. The mechanism of this phenomena was attributed to CCV in protein abundances, which were previously shown to correlate with mitochondria abundance [21]. However, this correlation is unlikely to be the entire story as a well known study demonstrated that the time to TRAIL-induced cell death of individual sister HeLa cells concomitantly treated with a potent inhibitor of protein translation, cyclohexamide, became less correlated with time [12]. As protein translation is inhibited, the cause for the depreciation of this correlation is unlikely to come from temporal fluctuations in protein abundance and may be attributed to the fluctuations in mitochondria abundances.

## Results

### Mitochondria density correlates with resistance to TRAIL

To assess whether mitochondria abundance correlated with single cell sensitivity to TRAIL induced apoptosis (Figure 1A) we measured the binary life-or-death status and the abundance of mitochondria of individual cells by flow cytometry. During extrinsic apoptosis, TRAIL stimulates cell death by binding to its cognate death receptors on the cell surface forming a complex that activates Caspase 8 (Figure 1A), the so-called initiator caspase (IC). Active IC in turn causes Bax accumulation and polymerization on the outer membrane of mitochondria, forming pores [22, 23] which allow for the diffusion of pro-apoptotic molecules from the inter-membrane space of the mitochondria into the cytosol [24, 25]. These molecules activate a cascade which ultimately induces the activity of Caspase 3, the so-called executioner (EC) caspase [24, 25], which is responsible for triggering cell death. In effect, these molecules dynamically regulate each other’s activity so that the continuous values of TRAIL concentration can be converted to a binary dead-or-alive response.

**Figure 1:**
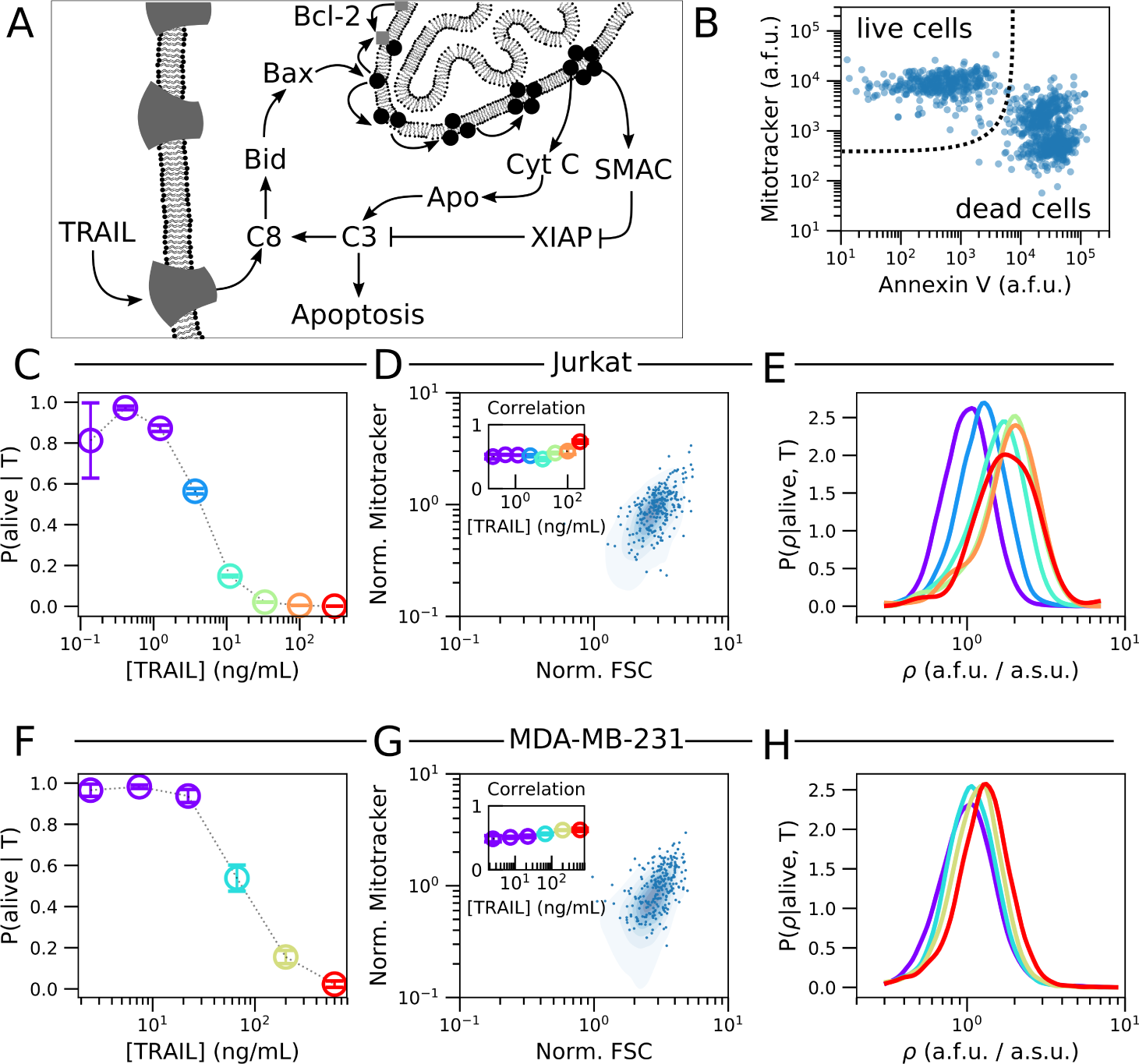
TRAIL administration enriches for cells with high density of mitochondria. (A) An overview of TRAIL induced apoptosis. (B) Flow cytometry measurements (FCM) of mitochondria (MitoTracker Deep Red) and phosphatidylserine (FITC conjugated Annexin V) in Jurkat cells. Complete flow cytometry gating strategy can be seen in Supplementary Figure 1. The fractional response of Jurkat cells (C) to TRAIL. Each color corresponds to a unique fractional response to a specific TRAIL dose. Cell size measurements (FSC-A) in Jurkat cells (D) are correlated with mitochondria abundance (MitoTracker Deep Red). The inset shows that the Pearson correlation marginally changes for each TRAIL dose. The probability density of mitochondria density (*ρ*) for each dose of TRAIL that elicits a unique response in Jurkat cells (E). The fractional response of MDA-MB-231 cells to TRAIL (F). Cell size measurements (FSC-A) in MDA-MB-231 cell (G) are correlated with mitochondria abundance (MitoTracker Deep Red). The inset shows that the Pearson correlation marginally changes for each TRAIL dose. The probability density of mitochondria density (*ρ*) for each dose of TRAIL that elicits a unique response in MDA-MB-231 cells (H). In (E) and (H) the single cell measurements from each of the lowest three doses of TRAIL are aggregated prior to probability density estimation (Violet). Visual inspection of the respective dose response curves suggest that these three doses of TRAIL are effectively identical. Data presented with errorbars represent the mean *±* one standard error of the mean over triplicate experiments.

The human T-lymphoblastoid derived cells (Jurkat), a human breast adenocarcinoma cell line (MDA-MB-231), and HeLa cells were exposed to different doses of TRAIL for four hours, a time frame in which cells died readily but the single cell abundances per mitochondria remained largely unchanged (Supplementary Figures 1 and 2). For each dose of TRAIL we measured the abundance of mitochondria and the cell state in single cells by concomitant labeling with a fluorescent Annexin V and MitoTracker Deep Red in flow cytometry measurements (FCM). Living cells, Annexin V negative and MitoTracker high, are well separated from the dead cells, Annexin V positive and MitoTracker low and medium (Figure 1B, see Supplementary Figure 1 for complete gating strategy). Importantly, the fact that the living and apoptotic cell populations shared no MitoTracker population lead us to conclude that the apoptosis process corrupted MitoTracker signal. Consequently, the apoptosis process precludes assessment of mitochondria abundance by MitoTracker in Annexin V positive cells.

From the FCM and our live cell gate we confirmed that Jurkat and MDA-MB-231 cell lines were sensitive to TRAIL (Figures 1C,F) but HeLa cells were not as responsive (see Supplementary Figure 9). Furthermore, from fitting the Hill model to each dose response we found that these cell lines had vastly different sensitivities (IC_50_) to TRAIL, 3.81*±*0.26 ng/mL for Jurkat cells, 76.4*±*8.77 ng/mL for MDA-MAB-231 cells and more than 300 ng/mL for HeLa cells. Because of this observation, we color-coded the effective abundance of TRAIL dose so that we may track the mitochondria abundance with the effective, as opposed to the actual, dose of TRAIL (Figures 1C-H and Supplementary Figure 9).

Next we found that mitochondria abundance of living cells is correlated with cell size, as measured by forward scatter (Figures 1D,G). To eliminate analyzing effects due to cell size, as opposed to mitochondria, we focus our attention to the mitochondria density, *ρ*, defined as the MitoTracker signal normalized to FSC signal. With these data we estimated the probability density of single cell mitochondria density in live cells for each dose of TRAIL. Here we find that with successively increasing doses of TRAIL the probability distribution *ρ* becomes increasingly enriched for cells with high mitochondria density (Figures 1E,H). Moreover, we find that the degree of the enrichment is unique to each cell line Jurkat cells are more readily biased in their mitochondria density than were the MDA-MB-231 and HeLa cells (for all HeLa cell analysis see Supplementary Note 6 and Supplementary Figure 9).

We hypothesize that the observed enrichment of cells with high mitochondria density is established by a differential sensitivity of single-cells to TRAIL. An intuitive result considering that the sensitivity of a signaling pathway to its cognate ligand is tuned by the abundances of its components. In apoptosis for example, we would expect that the number of TRAIL receptors on a cell’s surface, the number of pro-caspase molecules, the number of Bax molecules, the number of mitochondria, etc contribute to that cell’s response to a single dose of TRAIL. If each one of these molecules varied from one cell to the next, the so-called cell-to-cell variability (CCV), we should expect that the individual response of cells to TRAIL are unique.

Indeed, the probability density of *ρ* shows that the endogenous density of mitochondria vary from cell-to-cell (Figures 1E,H). If each cell’s sensitivity to TRAIL were anti-correlated with mitochondria abundance we would expect an enrichment of high mitochondria density cells with TRAIL stimulation. Such an effect can be quantitatively studied by using the rule of probability. By applying Bayes’ theorem we may associate the changes in the probability density *ρ* with the quantitative change of the fraction of living cells. From this simple property of probability, we were able to develop a quantitative strategy to gauge whether the observed endogenous variability of biological components are responsible for functional population diversity.

### Variability in all-or-none biological responses

As found in other biological systems, e.g. MAPK and NF*κ*B [26, 27], the conversion of a continuous input to a binary response limits the influence of CCV in cellular components to CCV in sensitivity to perturbations. In apoptosis, each cell, with its unique concentrations of molecular components, should require a specific concentration of TRAIL to induce cell death. At the population level the diversity in single-cell sensitivities to TRAIL gives rise to the fractional control of cell death.

As an example, consider two separate ensembles of cells, one with near identical biomolecular composition (low CCV) and the other with variable numbers of its components (high CCV). In the scenario where all components are near equal, the individuals will undergo the life-death transition at nearly the same dose of ligand (Figure 2A). In contrast, when CCV is relatively high, the individual cells of the ensemble will transition from live to dead at diverse doses of TRAIL (Figure 2B). The resulting fractional control of the population response to TRAIL would then take a steep or gradual sigmoid shape, respectively.

**Figure 2:**
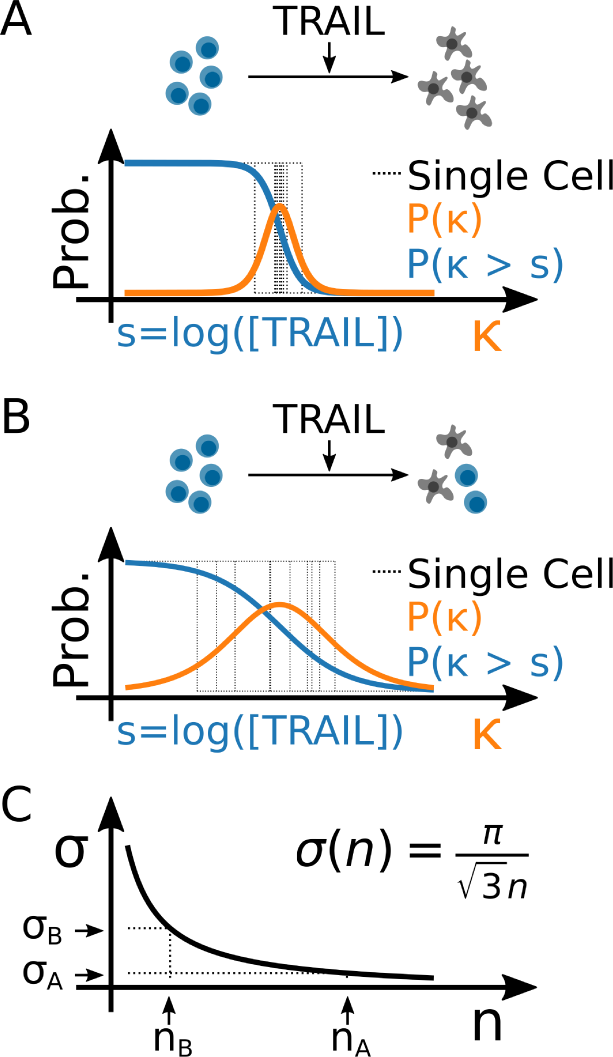
Cell-to-cell variability in the binary response to TRAIL. Hill response function with respect to TRAIL dose (blue) and the corresponding probability density of the single-cell sensitivities (orange) for populations with (A) low CCV and (B) high CCV. (C) The theoretical correspondence between the variance of single-cell sensitivities to TRAIL (*σ*) and the Hill coefficient *n*. Here, (*n_A_*, *σ_A_*) and (*n_B_*, *σ_B_*) represent the Hill coefficient and corresponding single-cell variances from (A) and (B), respectively.

This interpretation of the empirical dose response curve represents the cumulative distribution of single-cell sensitivities. From which, we may derive the corresponding probability density of single cell sensitivities. Indeed, from this simple interpretation, the empirical dose response curve of binary biological responses contains a complete statistical description of the functional diversity in the population. Fitting this dose response to a Hill function, we find that the mean sensitivity of single cells to a perturbation is simply the logarithm of the IC_50_, and the variance of single cell sensitivities to be inversely proportional to the squared Hill coefficient (Figure 2C).

Matching the dose response parameters to statistical quantities is useful, because now we may use the tools of probability theory to analyze our data such as taking conditional moments. In context of TRAIL induced apoptosis we may ask what is the average sensitivity of cells given a specific mitochondria density. This statistical question is equivalent then to asking how does the IC_50_ of individual cells change with the mitochondria density. Or we may ask, what is the variance of single cells sensitivities given that we measured mitochondria density. Which, intuitively, quantitatively measures the remaining diversity in the population once we remove the contribution of mitochondria density. With this information we may then compute the fraction of the functional relevant population diversity attributable to a measured component.

### Decomposing sources of cell-to-cell variability

Lets assume that the sensitivity of cells to TRAIL is wholly dependent on the biological components of the apoptotic signaling pathway. For simplicity lets designate the mitochondria density *ρ* to be *x*_0_ and all other contributing components as *x*_1_*, x*_2_*, &, x_m_*. *A priori* any mathematical function that describes the intricate relationships of these components and the dose of TRAIL to the single cell sensitivity (*κ*) is unknown, however we may expand this *a priori* unknown function to an arbitrary order by,

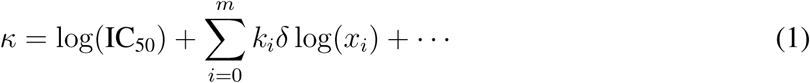

where *k_i_* = *∂κ/∂* log(*x_i_*)|_〈*xi*〉_ and *δ* log(*x_i_*) = log(*x_i_*) *−* log(〈*x_i_*〉) (see Supplementary Note 3 for details). The order in which we expand to will dictate the degree of complexity we wish to understand. If we limit our understanding to first order, then the details of the specific pathway are bundled into phenomenological parameters *k_i_*. If then, we infer *k_i_* from data, we can estimate the extent in which each component contributes to a cell’s sensitivity to TRAIL.

Indeed, Eq. 1 provides a framework for constructing a single cell interpretation of the Hill model, which incorporates the abundance of biological components with the stimulation strength. The biological species are introduced into the Hill model parameters by their influence on the first and second statistical moments of single cell sensitivities, *κ*. For example, incorporating our measurements of mitochondria density *ρ* to the IC_50_ amounts to computing the average sensitivity conditioned on mitochondria density,

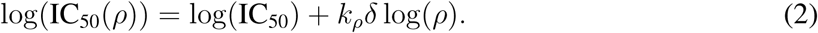

Then, in like fashion, the resulting Hill coefficient comes from estimating the variance of sensitivities conditioned on mitochondria density,

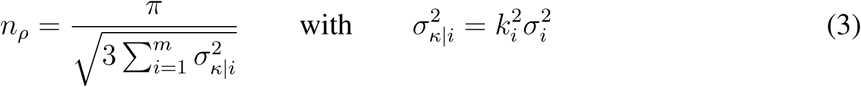

and 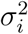 representing the variance of the *i^th^* biological component. If we apply the moments from our first-order expansion (Eqs. 2,3) to the Hill model we arrive at our single cell Hill model,

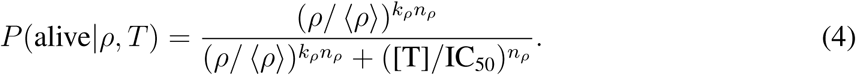

Eq. 4 gives us a detailed understanding on the influence of mitochondria density, or in general any measured components. If, for example, mitochondria density does not contribute to the cell’s sensitivity to TRAIL then *k_ρ_* = 0 and Eq. 4 reduces to the standard dose response Hill function. If however *k_ρ_* is not zero and positive, then mitochondria density effectively promotes cell survival, and if *k_ρ_* is negative then it increases the effectiveness of TRAIL. Together we can probe the influence of each measured component at unprecedented resolution. We call this strategy DEPICTIVE, which is an acronym for DEtermining Parameter Influence on Cell-to-cell variability Through the Inference of Variance Explained.

To see this in detail, lets consider an example of an arbitrary pathway consisting of three components that takes *s* as input and provides a binary output *y* (Figure 3A). We then make a synthetic data set representing virtual single cell flow cytometry measurements for different doses of *s* (Figure 3B). Using these data, we can compute the populations response to the stimulation, and due to the single cell nature of the data can interrogate the influence of each molecular constituent. The distribution of biological species *x* does not change whether we subset single cells upon their state *y* (Figure 3C). Consequently, *x* must not contribute to each cells sensitivity to *s*, a fact corroborated by *P* (*y* = 1*|x, s*) being invariant to the abundance of *x*. Unlike species *x*, species *z* and *q*do influence the cell’s behavior, which is apparent in analyzing the single cell data. Intuitively, the changes of the distribution of molecular components conditioned by cell state is the signal required for inferring each parameter *k_i_* from (Eq.1).

**Figure 3:**
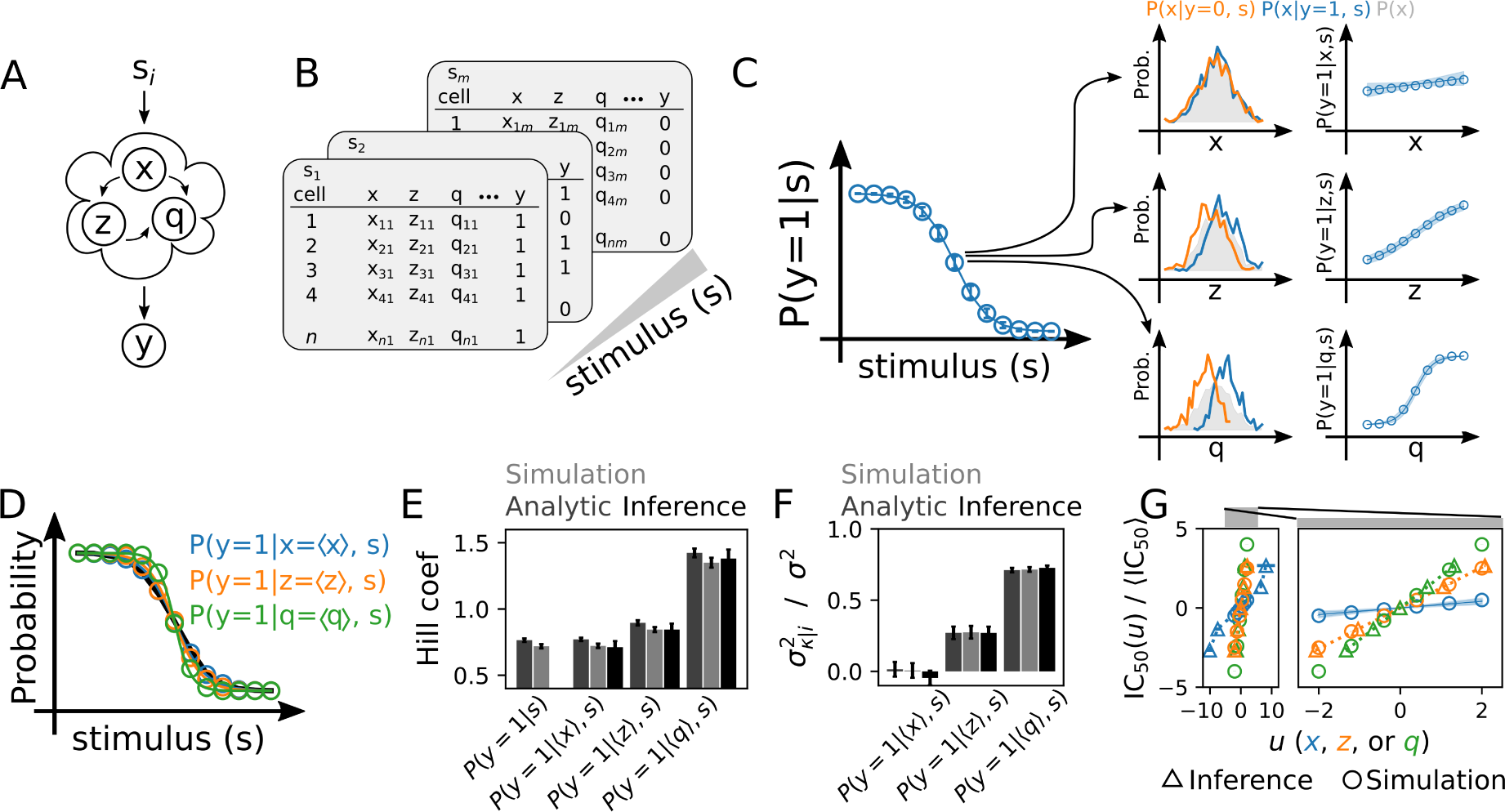
Decomposing sources of cell-to-cell variability. (A) Schema for a simple cellular response, *y* 0, 1, to the activation of pathway components *x, y, q* subject to the *i^th^* dose of a stimulus *s_i_*. (B) Single cell data were simulated to demonstrate the feasibility of CCV decomposition by sampling virtual cells (see Supplementary Note 3.2.3 for details). (C) The dose response of *N* = 1000 simulated cells and *n* = 100 replicate experiments in which *k_x_* = 0.25, *k_z_* = 1.25, and *k_q_* = 2. Error bars represent ± three standard deviations about the mean. The histograms normalized by cell count of live or dead cells reveals how each biological entity correlates with cell state (left column). We may make the dependence of cell survival to TRAIL by examining the probability of the cell state, *y* = 1, given the dose and abundance of each biological component (right column). In the right column, the circles represent the true conditional probability, while the blue line and shaded region represents the DEPICTIVE inferred dependence ± three standard deviations. (D) If we eliminate each individual source of CCV, the dose response is less uncertain. A phenomena that is well parameterized by the Hill coefficient (E), and the corresponding variance explained (F). Error bars represent ± one standard deviation. (G) The scaling of the IC_50_(*u*) with *u*.

Inferring each of the *k_i_* in the simulation data is trivial, because we have measurements of each biological component for each cell state *y*. Uniquely, our experimental data consists of MitoTracker measurements from live cells exclusively. This was because MitoTracker Deep Red signal is dependent on the electro-chemical properties of the mitochondria, which are different for live and dead cells. To infer the values of *k* from such data we developed a new inference strategy for semi-supervised logistic regression and embed it as module within the DEPICTIVE statistical framework (see Supplementary Note 3.3 for details). We apply our method to the synthetic data that is analogous to our measurements, that is the measurements associated with each virtual cell with a binary label *y* = 1. The filled regions of the curves in *P* (*y* = 1*|u, s*) for *u* = *x, z*, or *q* (Figure 3C) represent the model predictions from inferred *k_u_ ±* 3 standard deviations. These inferred parameters were then used to infer the single cell dose functions (Figure 3D) with qualitatively excellent agreement. Quantitatively, we see that we can infer the conditional Hill coefficients (Figure 3E), the corresponding variances explained (Figure 3F), and lastly the dependence of the single cell sensitivities on each biological component (Figure 3G).

### Mitochondria density is a source of cell-to-cell variability

We apply our new statistical framework, DEPICTIVE, to quantitatively dissect the dependence of single cell sensitivities to TRAIL with mitochondria density (Figures 1E,H). We see that the fractional response of the Jurkat cells to each dose of TRAIL, *P* (alive*|ρ, T*), is strongly dependent on mitochondria density (Figure 4A, see Supplementary Figures 3-5 for goodness-of-fit analysis). Moreover, we see that the single cell dose response curve translates from low TRAIL to high TRAIL doses with increasing mitochondria density (Figure 4B). The MDA-MB-231 cells fractional response (Figures 4C,D) is less steep than that of Jurkat, indicating that the single cell sensitivities of MDA-MB-231 cells to TRAIL are not as sensitive to CCV in mitochondria density as Jurkat (see Supplementary Figures 6-8 for goodness-of-fit analysis). A result that can be summarized by plotting the IC_50_(*ρ*) for each cell line (Figure 4E). Moreover, we find that 30% and 2% of the diversity in single-cell sensitivities to TRAIL may be attributed to mitochondria density in Jurkat and MDA-MB-231 cells, respectively (Figure 4F).

**Figure 4:**
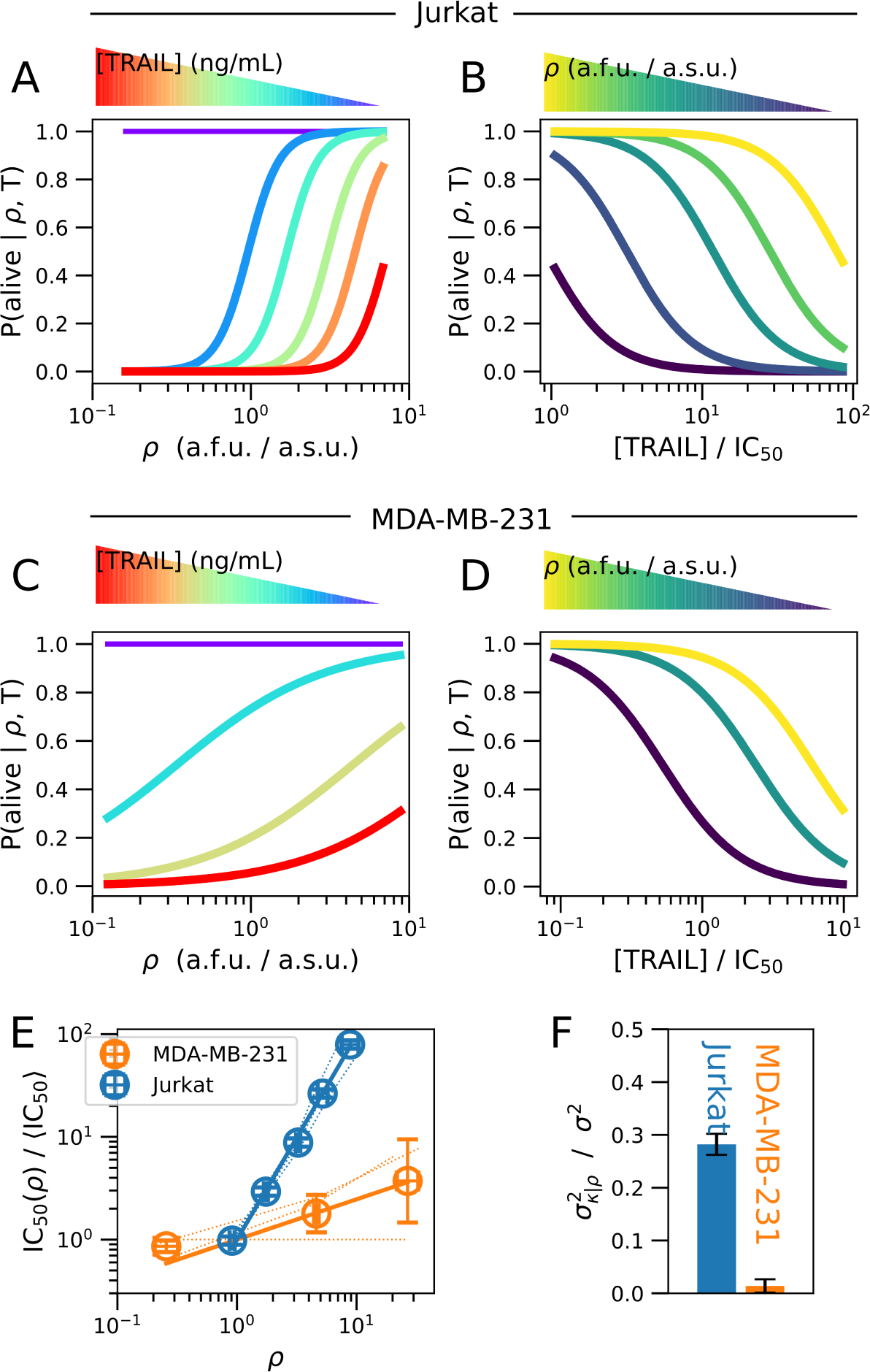
CCV in mitochondria density influences fractional response to TRAIL. The inferred fractional response of Jurkat cells (A) or an MDA-MB-231 cells (C) as a function of *ρ* given TRAIL dose. The inferred fractional response of Jurkat cells (B) or an MDA-MB-231 cells (D) as a function of TRAIL dose given *ρ*. (E) The dose of TRAIL normalized by the population IC_50_ (y-axis) and the inferred density of mitochondria (*ρ*) in which the fractional response is 0.5 (x-axis). The blue markers represent triplicate averages in Jurkat cells while the error bars represent the standard er ror of the mean. The orange markers represent duplicate averages in MDA-MB-231 cells and standard error of the mean, while the y-axis represents triplicate statistics. We report duplicate statistic in MDA-MB-231, because in one replicate experiment there is no correlation between the IC_50_ and *ρ*, including these values of *ρ* would lead to misleadingly large error bars. Lastly, dashed blue and orange lines represent the inferred values of *ρ* for each replicate data set for Jurkat and MDAMB-231 cells, respectively. (F) The fraction of the variance in single cell TRAIL sensitivities (*σ*) explained by CCV in mitochondria density from (*σ_κ|ρ_*). Error bars represent standard error of the mean of experimental triplicates. Detailed analysis of each replicate set are presented in Supplementary Figures 3-5.

### Bax concentration dependence on mitochondria surface area

To gain mechanistic insight in the functional role of mitochondria density in the cell death decision, we developed a coarse-grained dynamic model of apoptosis (Figure 5A). Our description aims to reproduce the dominant dynamical features of initiator caspase reporter protein (IC-RP) first measured and published by [24]. These being a slow but accelerating initial increase of IC followed by a fast increase in both IC and EC. To such end our model includes: i) a slow auto-catalytic increase in IC activation, ii) a quasi-steady-state approximation for Bax pore formation dynamics and mitochondrial outer membrane permeabilization (MOMP), and iii) the strong positive feedback from EC to IC (see Supplementary Note 5 for details).

**Figure 5:**
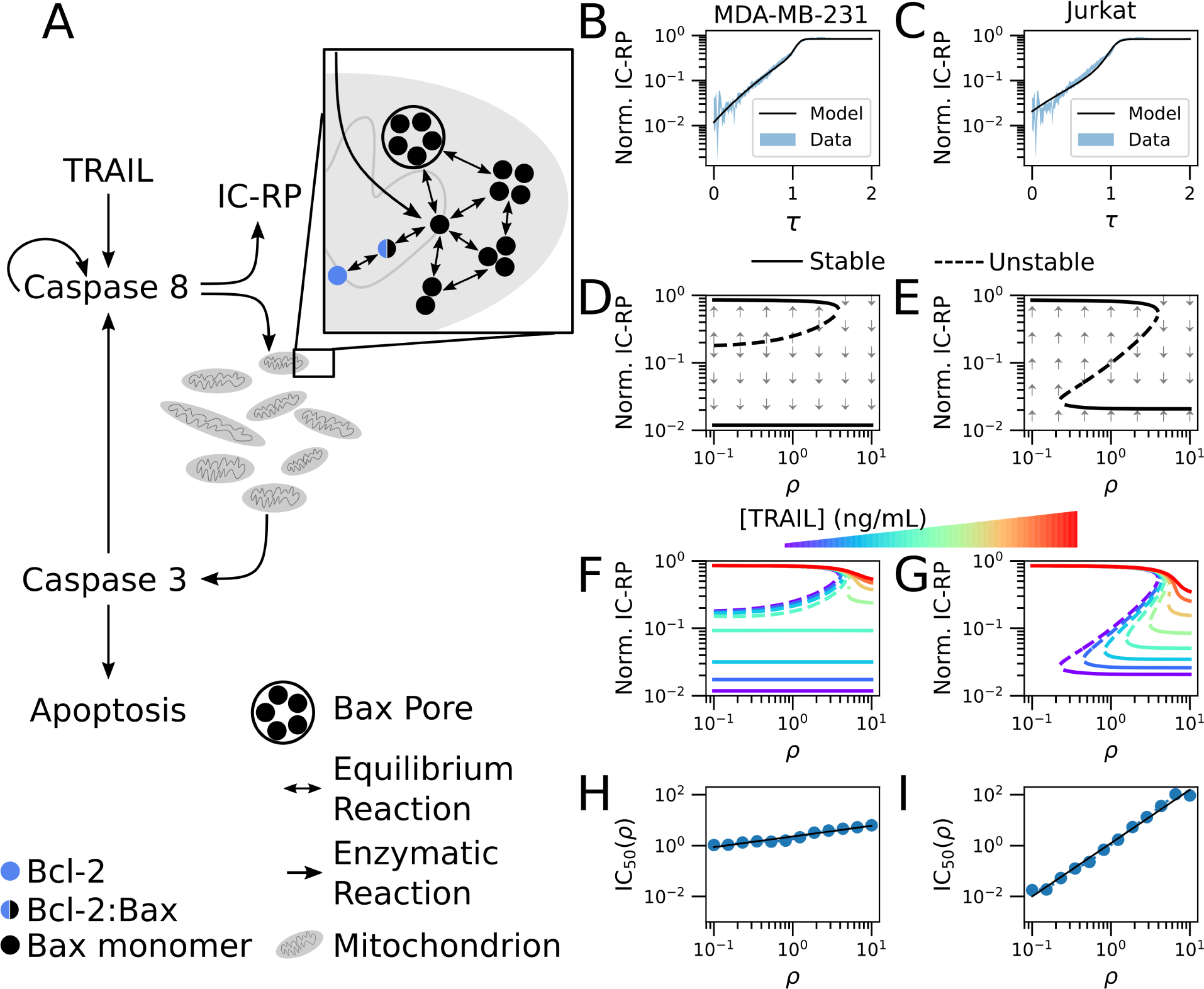
Mechanism of IC_50_ dependence on mitochondria density. (A) Simple model of apoptosis. The dynamics of initiator caspase reporter protein (IC-RP) from [24] and the model-inferred dynamics corresponding to (B) MDA-MB-231 and (C) Jurkat cell lines. The model bifurcation diagrams for [TRAIL] = 0 ng/mL in (D) MDA-MB-231 and (E) Jurkat cells. The influence of TRAIL dose on the model fixed points for (F) MDA-MB-231 and (G) Jurkat cells. The dependence of single cell sensitivities to TRAIL on *ρ* for (H) MDA-MB-231 and (I) Jurkat cells. The IC_50_ was estimated from Hill function fits of simulated data (blue circles), and which were then fit to a power law (black line). Simulations consisted of 100 cells per each of the 20 doses of TRAIL and 12 densities of mitochondria considered.

We conjectured that TRAIL induced activation of IC in Jurkat and MDA-MB-231 cells match the biphasic increase of IC-RP measured in HeLa cells (Albeck et al. 2008b), but differ in their propensity to form Bax pores (Figures 5B,C). Specifically, we consider unique susceptibilities of Bax pore formation to Bcl-2 mediated inhibition for each cell line. As Bax pores reside in the mitochondria, the effective Bax concentration for a given amount of Bax decreases with mitochondria density. Implementing this insight into the model equations we see that the influence of mitochondria density can be understood through the corresponding bifurcation diagrams (Figures 5D,E).

The dynamic properties of IC in MDA-MB-231 cells in the absence of TRAIL are either bistable or monostable depending on mitochondria density. In these diagrams, the high IC fixed point corresponds to cells that have integrated sufficient signal for MOMP and consequently represent apoptotic cells. Cells with relatively low mitochondria density are bistable and may undergo apoptosis only if their IC abundance exceeds a critical amount designated by the dashed line (Figure 5D). This bistable region does not preclude cell death cells may acquire sufficient abundances of IC for death by fluctuations in biomolecular reactions. Indeed, the likelihood of such an event decreases with the difference of IC abundance between the unstable fixed point (dashed line) and low IC stable fixed point (solid line). Meanwhile, cells with relatively high mitochondria density only have a single fixed point of low IC, indicating that these cells will never spontaneously undergo apoptosis in the absence of TRAIL.

In contrast, the bifurcation diagram representing Jurkat cells shows three distinct regions (Figure 5E). These being cells with: 1) low density of mitochondria having a single fixed point of high IC and consequently all die; 2) medium density of mitochondria are bistable, for which, the fractional response to TRAIL decreases with the concomitant increase in the IC unstable fixed point and mitochondria density; and 3) high density of mitochondria are monostable with low IC abundances, hence all cells survive. Next, we extend these analyses to the full range of TRAIL doses.

The influence of increasing TRAIL dose in each cell type specific parameterized model is evident in their bifurcation diagrams. MDA-MB-231 cells respond to TRAIL by increasing the IC abundance of the lower fixed point (Figure 5F). In doing so, cells with mitochondria density in the bistable region equally increase their susceptibility to cell death from fluctuations in IC abundance. The Jurkat model’s response to TRAIL exhibits an increase of the density of mitochondria that separates the monostable high and bistable IC abundance regions (Figure 5G). Therefore, an individual cell’s mitochondria density determines its sensitivity to TRAIL induced cell death. Together, these model-based observations propose an explanation for how CCV in mitochondria density influences the response of Jurkat but to a lesser extent MDA-MB-231 cells to TRAIL (Figures 5H,I).

### Sensitizing MDA-MB-231 cells to CCV in mitochondria density

In inspecting the model parameters associated with each cell type, we noticed that MDA-MB-231 cells were more susceptible to Bcl-2 mediated inhibition of Bax pore formation than Jurkat. We hypothesized that this effect would be abated by incorporating a small molecule inhibitor to Bcl-2 in MDA-MB-231 cells (Figure 6A, see Supplementary Note 5.1 for derivation). By incorporating Bcl-2 inhibition, we found that the sensitivity of the fractional response of the cell population to TRAIL increases (Figure 6B). Furthermore, and as intuited, Bcl-2 inhibition increased the dependence of single-cell sensitivities to TRAIL on mitochondria density (Figure 6D). We corroborated these theoretical predictions by measuring the influence of the clinically relevant small molecule inhibitor of Bcl-2 family proteins ABT-263 [28] (Figures 6C,E, see Supplementary Figures 6-8 for goodness-of-fit analysis). Remarkably, Bcl-2 inhibition alone increased the variance of sensitivities attributable to mitochondria density from 0% to 40% (Figure 6F).

**Figure 6:**
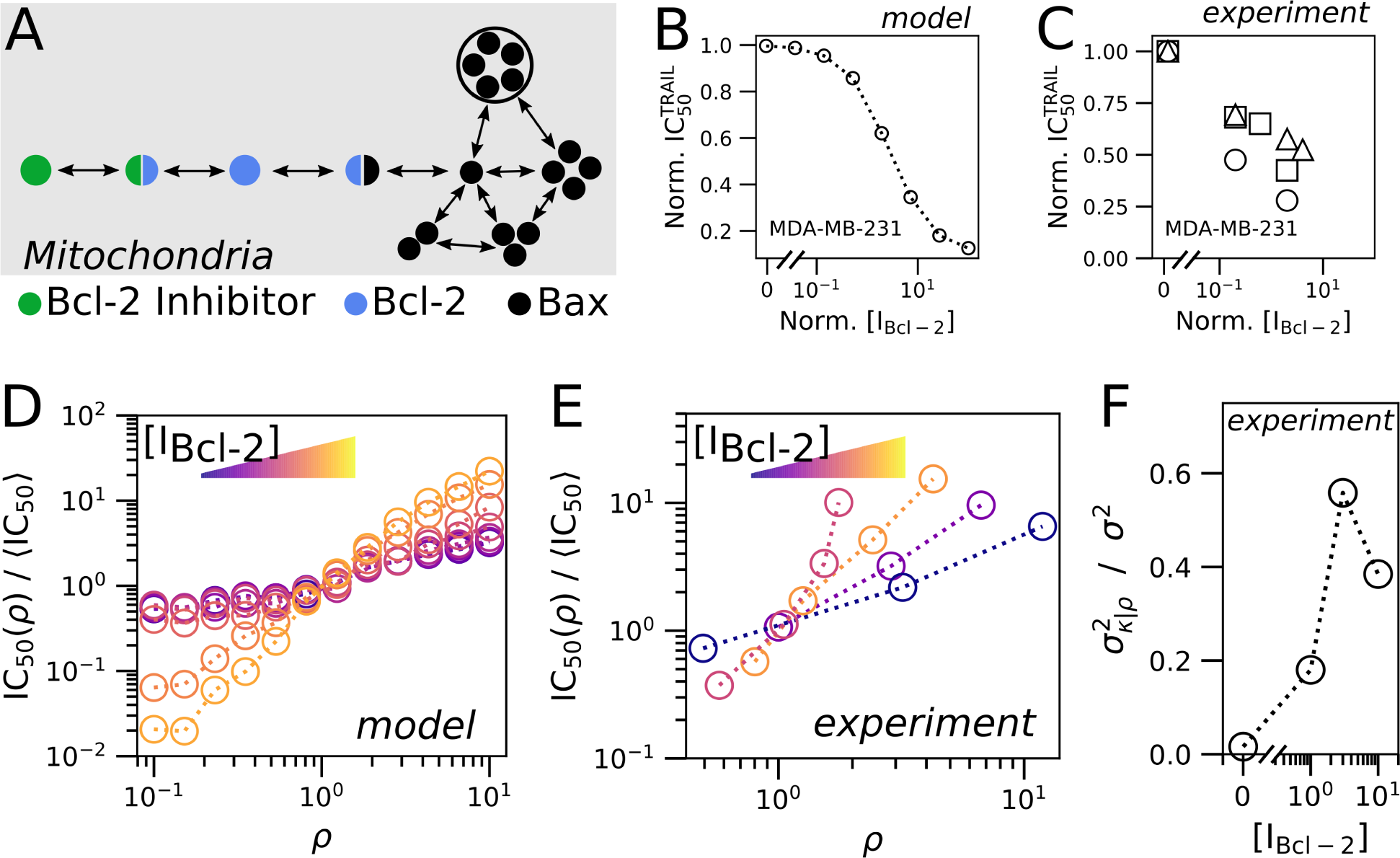
Plasticity in fractional response to TRAIL. (A) Bcl-2 inhibitor reduces the effective abundance of Bcl-2 by formation of Bcl-2:Bcl-2 inhibitor complex. (B) Simulation results of the population IC_50_^TRAIL^ response to Bcl-2 inhibition in MDA-MB-231 cells. (C) Experimental measurement sets uniquely represented by a square, circle or triangle marker of the population IC_50_^TRAIL^ response to Bcl-2 inhibition for MDA-MB-231 cells. (D) Estimated IC_50_ for changing *ρ* from MDA-MB-231 parameterized model simulations. (E) The experimental dependence of IC_50_ on *ρ*, from a single representative experiment of three replicate experiments (Supplmenetary Figures 6-8), as computed in Figure 4E for [0, 1, 3, 10] *µM* doses of the Bcl-2 Inhibitor ABT-263. (F) The fraction of variance in single-cell sensitivities (*σ*) explained by mitochondria density CCV in E (*σ_ρ_*). Note that all simulations were conducted with 100 cells for each of the 20 doses of TRAIL, 12 densities of mitochondria, and 9 doses of inhibitor. Detailed analysis of each replicate data set are presented in Supplementary Figures 6-8.

## Discussion

We have unveiled a connection in the cell-to-cell variability of mitochondria density to the fractional control of TRAIL induced cell death. Importantly, we find that the dependency of single cell sensitivities to CCV in mitochondria abundance is cell line dependent. Presumably this dependence originates in the unique composition of components across cell lines. In that, the functional manifestation of CCV in mitochondria on the sensitivity of single cells to TRAIL induced apoptosis is dependent on the relative abundance and diversity of mitochondria in relation to the other biological constituents in the apoptosis pathway. Indeed, Jurkat cells readily responded to TRAIL and its IC_50_ scaled with mitochondria abundance, MDA-MB-231 cells showed scaling and responded readily to TRAIL only in the presence of a pan-Bcl-2 inhibitor, while scaling was never observed and only a minority of HeLa cells responded to TRAIL even during Bcl-2 inhibition. Consequently, the seemingly contradictory results of our study and that of Márquez-Jurado [20] are manifestations of the unique biological systems being studied. In particular, we think that our observations are highlighting a different phenomenon than Márquez-Jurado et al. who are measuring a mitochondrial dependence of cell death in HeLa cells regardless of TRAIL dose (see Supplementary Note 6 and Supplementary Figure 9).

Our findings were established by a new statistical framework, DEtermining Parameter Influence on Cell-to-cell variability Through the Inference of Variance Explained, namely DEPICTIVE, we developed to measure the impact of CCV on the binary response of cells to perturbation. It is composed of two parts, the first part is to infer the parameters of the logistic regression model when data from one or both of the binary cell state labels are available. While the second part provides the mathematical bases for interpreting the logistic regression model parameters to compute useful quantities.

Indeed, inferring the parameters of a logistic regression model from data is commonplace. However, it is only commonplace when data representative of both of the corresponding binary states is well established. To our knowledge, there is no method to infer these parameters from data where only one of the binary classes is readily available. In our study data from live and dead cells were unavailable because our experimental label of mitochondria abundance, MitoTracker DeepRed, was not reliable for dead cells.

The second part of DEPICTIVE statistical framework is to use the logistic model parameters to estimate the contribution of the measured biological component(s) to the variable binary response of single cells. Applying this tool, we found that mitochondria density accounts for nearly 30% of the variable response to TRAIL in Jurkat cells and varies from 2% to up to 40% in MDA-MB-231 cells when Bcl-2 is inhibited. Conversely HeLa cells showed no mitochondrial density dependence. Together, the two parts of the DEPICTIVE statistical framework can extract quantitative insights to sources of cell-to-cell variability.

We attribute the measured connection of TRAIL sensitivity and mitochondria density to the dilution of Bax on the outer mitochondrial membrane in cells by mathematical modeling. From the quantitative insights of DEPICTIVE, we found that the functional manifestation of mitochondrial CCV is plastic readily and predictably tunable by small molecule inhibitors of Bcl-2. It is plausible that this plasticity is a tool accessible to cells, and therefore may be co-opted by pathological cellular populations. For example, high mitochondria abundance can be a non-genetic mechanism of resistance to pro-apoptotic therapeutics. Incorporation of such knowledge may be an important consideration in developing therapeutic strategies.

The observed advantage of cells with high mitochondria densities may manifest in time-scales much longer than the life span of a single cell or the disease in a human, but propagate to the long time-scales of evolution. To date, the evolutionary hypothesis of mitochondria is as a symbiotic bacterium inside a proto-eukaryotic cell [29], exchanging safety for energy. However, another such evolutionary advantage may be expected, that this symbiosis would create a survival advantage such as the one described here. These results suggest that environmental constraints can select subpopulations not only based on genetic composition, protein abundances, but also CCV in organelle abundances.

## Methods

### Cell culture

**Jurkat** E6-1 cells originate from a male human acute T cell Leukemia and were purchased from ATCC (TIP-152). Cells were cultured in RPMI-1640 medium (Corning cat. 10-040-CV) supplemented with 10% heat inactivated fetal bovine serum (Corning cat. 35-011-CV), 2mM LGlutamine (Corning cat. 25-005-CI) and 1mM sodium pyruvate (Corning cat. 25-000-CI). Cells were cultured at 37^*◦*^ C in 5% CO_2_ in a humidified incubator and maintained at cell density not exceeding 3 x 10^6^ by addition of fresh medium, or by centrifugation with subsequent resuspension at 1 x 10^5^ cells/mL.

**MDA-MB-231** cells originate from a human female adenocarcinoma that were harvested from a metastatic cite in the breast. Cells were cultured in DMEM medium (Corning cat. 10-017-CV) supplemented with 10% fetal bovine serum and 2mM L-Glutamine (Corning cat. 25-005-CI). Cell were cultured at 37^*◦*^ C in 5% CO2 in a humidified incubator and subcultured every 2-3 days with 0.25% trypsin (Corning cat. 25-053-CI) to maintain sub-confluent density.

**HeLa** cells were purchased from ATCC (ATCC CCL2). Cells were cultured in DMEM medium (Corning cat. 10-017-CV) supplemented with 10% fetal bovine serum and 2mM L-Glutamine (Corning cat. 25-005-CI). Cell were cultured at 37^*◦*^ C in 5% CO2 in a humidified incubator and subcultured every 2-3 days with 0.25% trypsin (Corning cat. 25-053-CI) to maintain sub-confluent density.

## Apoptosis assay and Data acquisition

**Jurkat cells** were pelleted by centrifugation for 5 minutes at 100 x g, and then resuspended in 1x PBS and stained with 200 nM MitoTracker Deep Red (Life Technologies, cat. M22426) for 10 minutes at 37^*◦*^ C. MitoTracker staining was quenched with full cell culture medium, followed by centrifugation for 5 minutes at 100 x g. Cells were resuspended in cell culture media at a density of 1 x 10^6^ per mL, in which 1 x 10^5^ were transferred to each experimental well of a flat-bottom 96-well plate. Cells were then incubated at 37^*◦*^ C for 4 hours with different doses of Superkiller TRAIL (Enzo Life Sciences cat. ALX-201-115) and/or ABT263 (ApexBio cat. A3007). After drug treatment, cells were transferred to a v-bottom 96-well plate, pelleted by centrifugation at 1,000 x g, stained with FITC-conjugated Annexin V (Biolegend cat. 640945), and then measured by flow cytometry.

**MDA-MB-231 or HeLa** cells were seeded on 12-well plates at 5 x 10^5^ cells per well in 400 *µ*L, incubated overnight at 37 C in 5% CO2 in a humidified incubator until 80% confluent. Cells were then washed once with PBS and stained with 200 nM MitoTracker Deep Red (Life Technologies, cat. M22426) for 10 minutes at 37 C. MitoTracker staining was quenched with full cell culture medium, and then incubated at 37 C for 4 hours with different doses of Superkiller TRAIL (Enzo Life Sciences cat. ALX-201-115) and/or ABT263 (ApexBio cat. A3007). After drug treatment, supernatant containing floating cells was collected, and the remaining adherent cells were trypsinized, pooled with the supernatant, and pelleted by centrifugation for 5 minutes at 1,000 x g. Cells were then stained with FITC-conjugated Annexin V (Biolegend cat. 640945), and then measured by flow cytometry.

Flow cytometry measurements were conducted on a BD LSRII mainted by the Icahn School of Medicine at Mount Sinai flow cytometry core facility.

## FCM gating

FCM measurements were gated as follows: to exclude debris (Supplementary Figure 1A), then gated for singlets (Supplementary Figure 1B), MitoTracker Deep Red positive (Supplementary Figure 1C), and lastly for living cells by Annexin V (Supplementary Figure 1D). The fraction of cells alive was computed by dividing the number of cells in the Annexin-V-negative gate by the number of cells of the MitoTracker Deep Red positive gate. Subsequent single cell analysis was then conducted exclusively using cells from the Annexin-V-negative gate.

## Code availability

### DEPICTIVE

Detailed derivation of the DEPICTIVE strategy can be found in Supplementary Note 3. We developed a user friendly Python package to run the DEPICTIVE analysis strategy. The code is freely available as a GitHub repository, https://github.com/robert-vogel/depictive. Along with these tools we provide two tutorials that demonstrates how to generate synthetic data and to apply DEPICTIVE analysis. These tutorials can be found on the repositories wiki pages, https://github.com/robert-vogel/depictive/wiki.

### Dynamics Simulations

Detailed derivations and parameter values of model equations for simulation can be found in Supplementary Note 5. We developed a user friendly Python package to run, plot, and perform basic analysis of our model. The code is freely available as a GitHub repository, http://github.com/robert-vogel/mito_sims. Along with these tools we provide a a series of tutorials that demonstrates the use of our tools by examples. These tutorials can be found on the repositories wiki pages, https://github.com/robert-vogel/mito_sims/wiki.

## Data availability

The data presented in the main-text of this paper can be found on Mendeley data [30–35].

## Modeling and Statistical analysis

Detailed derivations of our DEPICTIVE statistical framework, application of DEPICTIVE to data, dynamics models, and inference of dynamic model parameters can be found in Supplementary Notes 3, 4, 5, and 5.4, respectively.

## Acknowledgments

This research was funded in part by grants from the NIH, (P50GM071558 (Systems Biology Center New York), R01GM104184 and U54HG008098 (LINCS Center) for M.R.B., R01 CA206005 for J.E.C. and P30 CA196521 a Tisch Cancer Institute Cancer Center Support Grant) and also an IBM faculty award for M.R.B. We wish to thank John Albeck for sharing single cell data and staff of the flow cytometry core facility at ISMMS and Julia Moore Vogel for help with the manuscript.

## Author contributions

Conceptualization, L.C.S., R.V., M.R.B., G.S., and P.M.; Methodology, R.V., G.S., and P.M.; Software, R.V., and P.M.; Formal Analysis, R.V., and G.S.; Investigation, L.C.S., R.V., and P.M.; Resources, J.E.C., M.R.B., and G.S.; Writing Original Draft, L.C.S, R.V., and P.M.; Writing Review & Editing, L.C.S., R.V., M.R.B, G.S., and P.M.; Visualization, R.V.; Supervision, M.R.B., G.S., and P.M.

## Materials and Correspondence

Requests for data, resources, and or reagents should be directed to Pablo Meyer.

